## Supplementary Information for "Subcortical correlates of consciousness with human single neuron recordings"

### Supplemental information

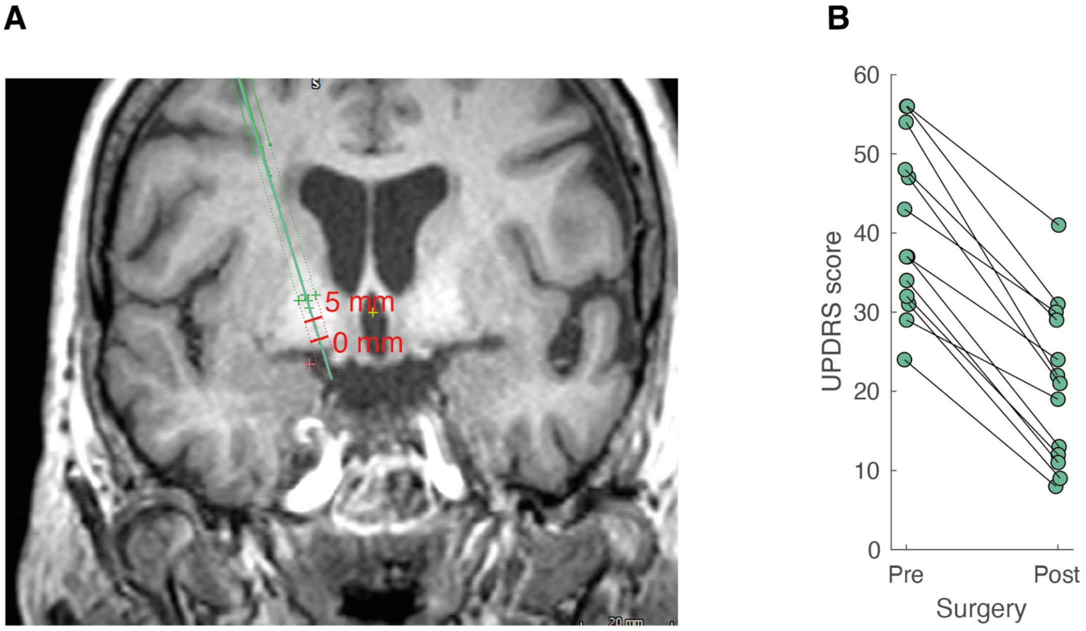


***Figure S1. Confirmatory localization analysis. A***. *Tract reconstruction based on pre-surgery MRI and post-surgery CT images (green segment) with red markers at 5 mm above and at target (0 mm) corresponding to depths in Figure 2.* ***B***. *Improvement in UPDRS scores pre- and post-surgery.*

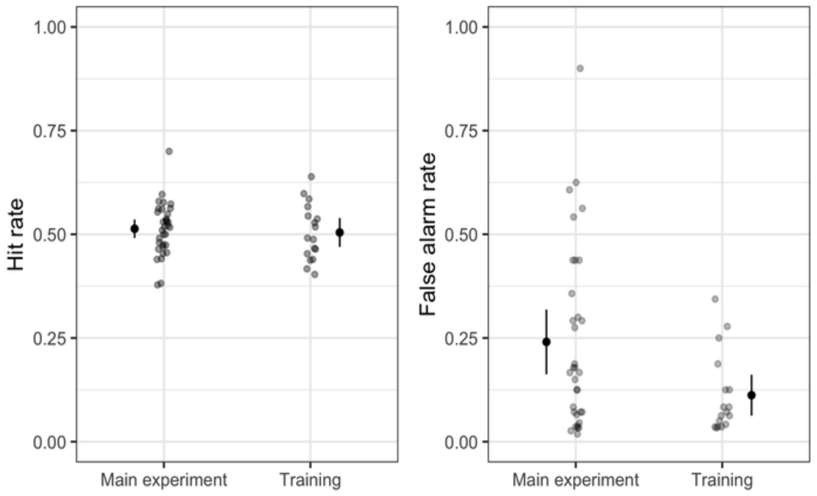


***Figure S2. Hit rate and false alarm rate observed during the main experiment and the training session.*** *Each small dot represents a participant. Big dots represent averages; error bars represent 95% confidence intervals.*

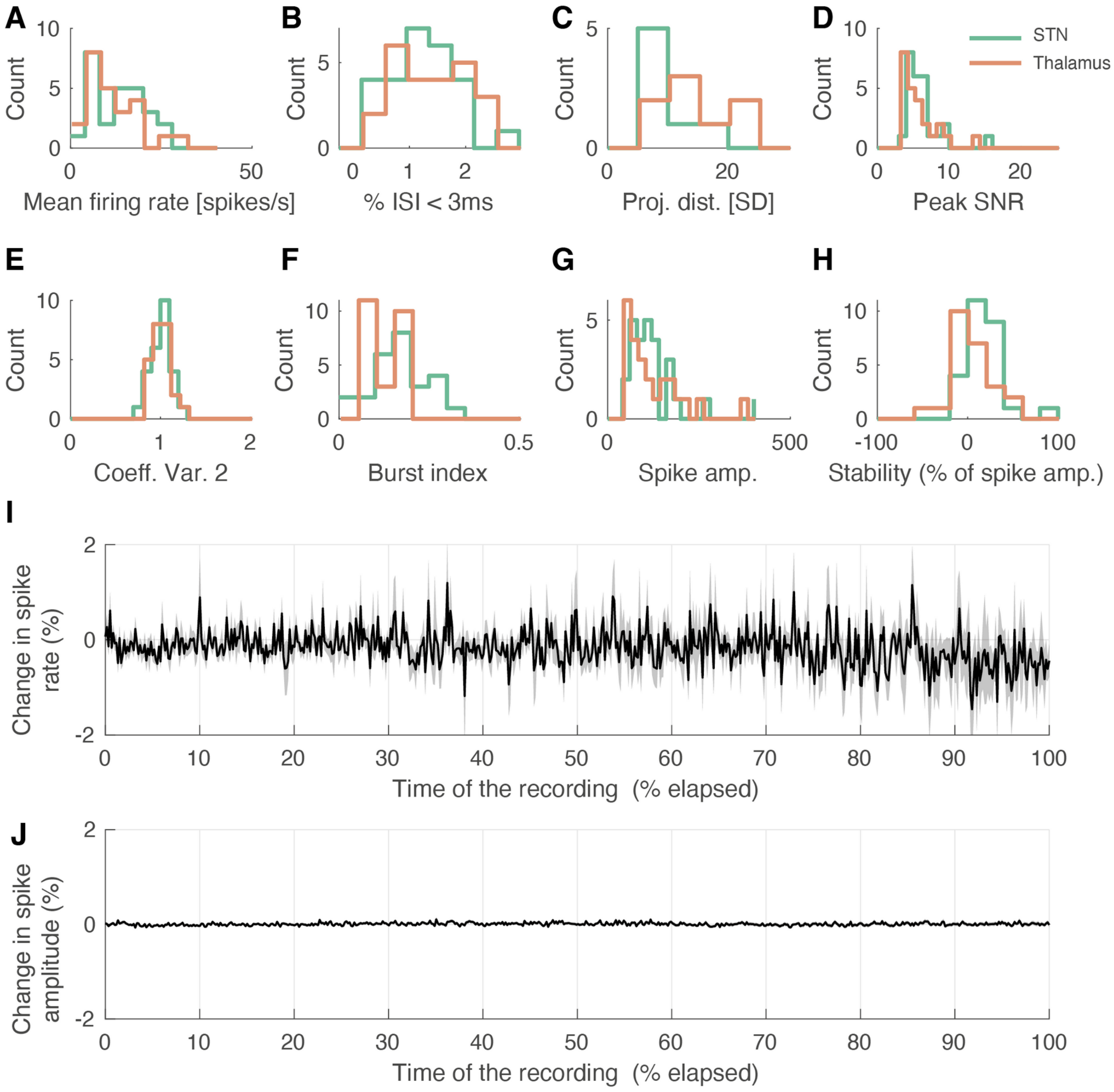


***Figure S3. Spike sorting metrics***. ***A***. *Histogram of firing rate during the time recordings were analyzed for neurons in the STN (green) and thalamus (orange).* ***B***. *Percentage of inter-spike intervals falling within a 3 ms refractory period.* ***C***. *Projection distance in standard deviation (SD).* ***D***. *Peak signal-to-noise ratio (SNR). E. Modified coefficient of variation (Coeff. Var. 2).* ***F***. *Burst index (BI).* ***G***. *Average spike amplitude in μV.* ***H***. *Percentage of change in spike amplitude between the beginning and end of time recordings were analyzed.* ***I***. *Change (in percent related to start) in the firing rate reinterpolated for comparison across sessions.* ***J***. *Change (in percent related to start) in the spike amplitude, reinterpolated for comparison across sessions. The shaded area represents the bootstrapped standard error of the mean across recordings.*

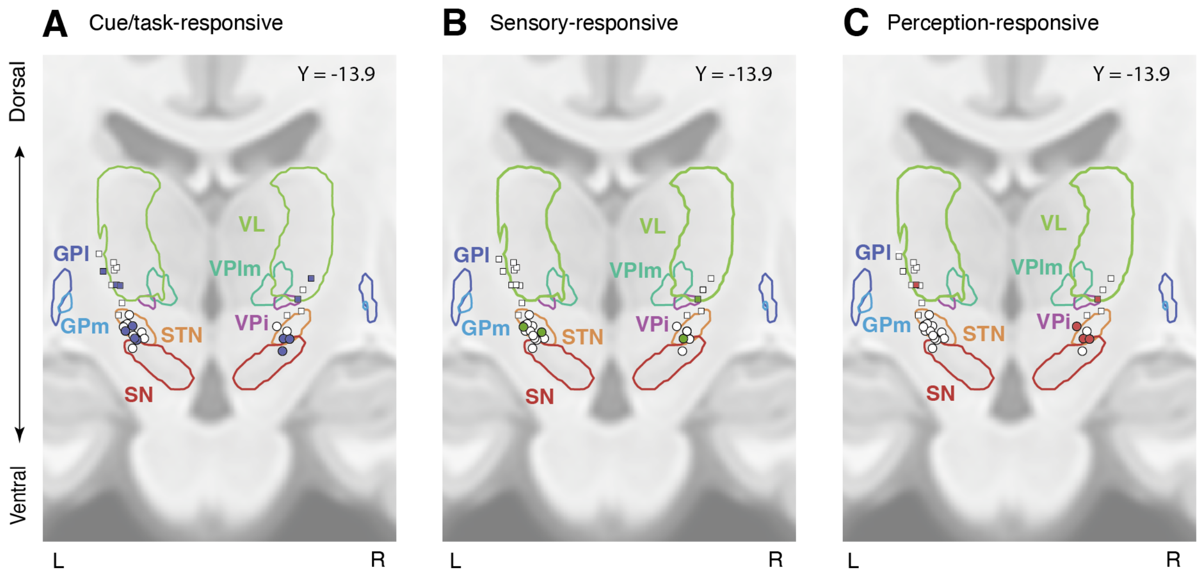


***Figure S4. Coronal view of recording locations.*** *Thalamic (squares) and subthalamic (circles) targets (see Figures 2-4 for a sagittal view) for patients for which we could obtain anatomical images . Filled circles or squares correspond to **A**. Task-responsive **B**. Sensory-responsive, or **C**. perception-selective neurons. Legend: VL: ventral lateral thalamus, VPlm: ventral posterior lateral and medial thalamus, VPi: ventral posterior inferior thalamus, STN: subthalamic nucleus, SN: substentia GPi/e: globus pallidus internalis / externalis,*

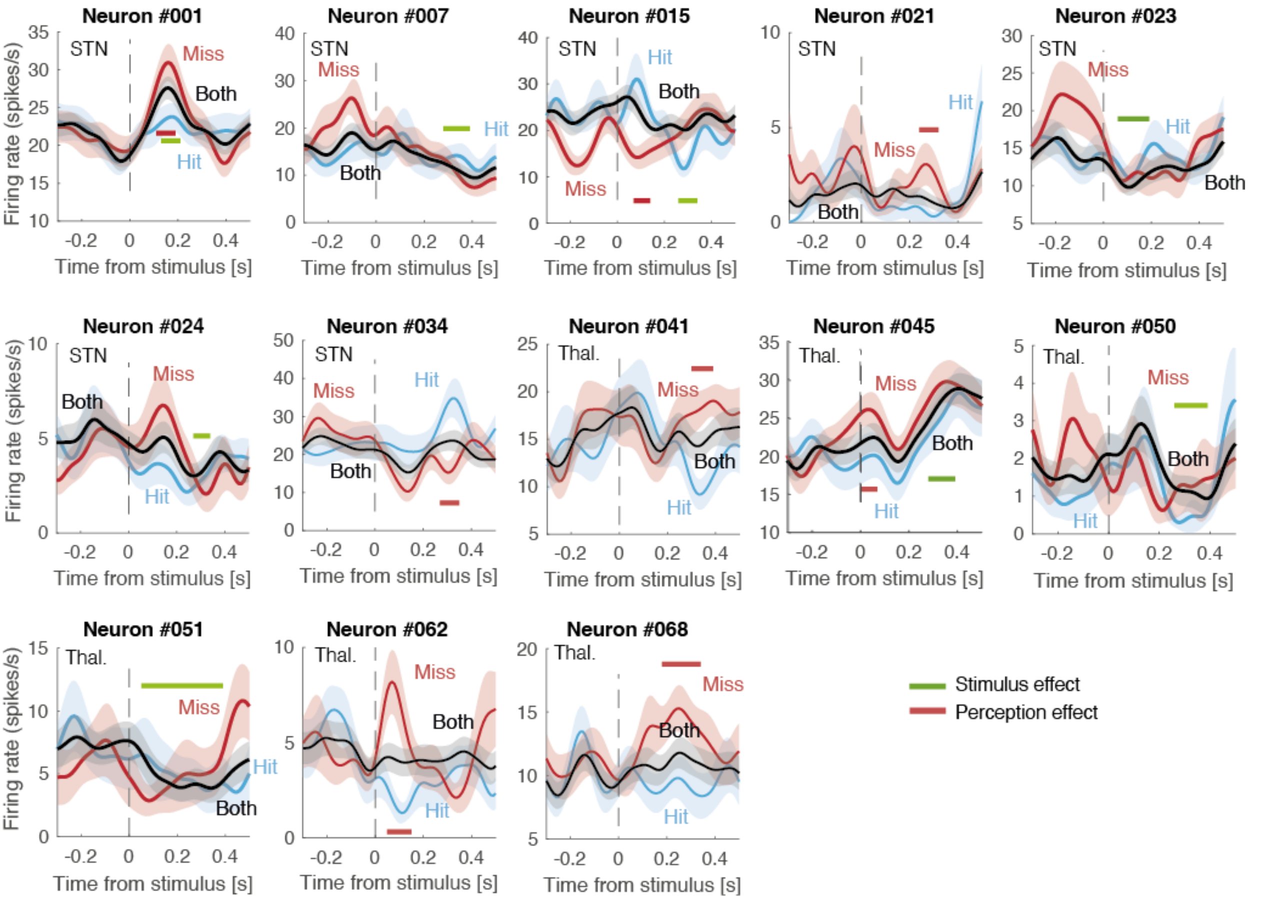


***Figure S5. Firing rate time-locked to the onset of the stimulus** (100 ms vibrotactile stimulation; vertical dashed line) for hits (light blue), misses (red) and for combined data (both; black). Thick horizontal segments show significant time windows for stimulus- (green) and perception-selective neurons (red). Shaded areas indicate bootstrapped standard errors.*

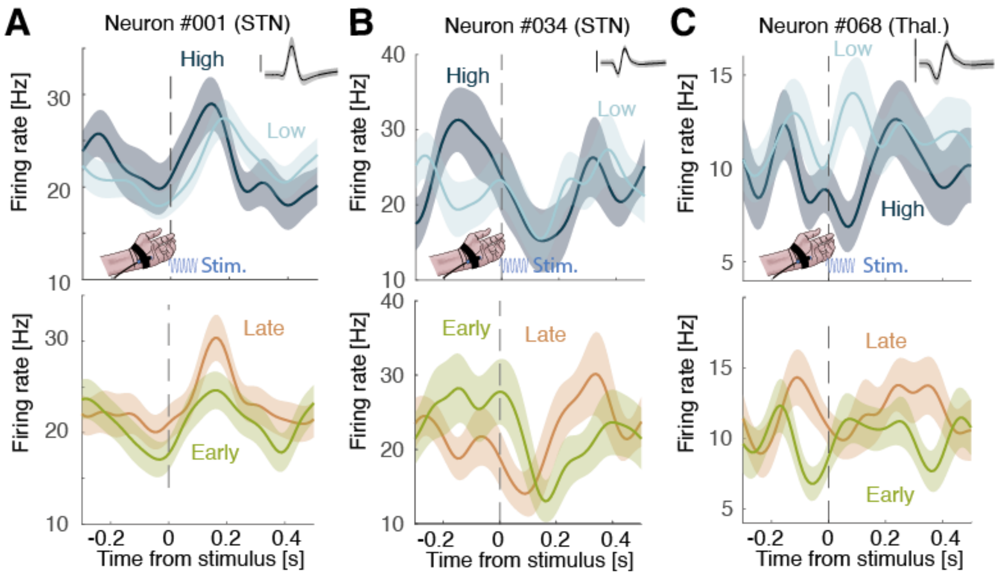


***Figure S6. Neurons from Figure 4, for different stimulus intensities and onsets.*** *We used the same trials as in Figure 4 but segregated in high versus low stimulus intensities (upper panel) or short and long stimulus onsets (lower panel). We found only 5 / 32 neurons sensitive to stimulus intensity (16%; p = 0.13; permutation test) and no neurons sensitive to stimulus onset (0 / 35). None of the 5 intensity-selective neuron corresponded to a perception-sensitive neuron.* ***A-C***. *Firing rate time-locked to the onset of the stimulus (100 ms vibrotactile stimulation; blue sinusoid) for high intensity (light blue) and low intensity (dark blue) trials (upper panel) or early (green) and late (orange) stimulus onsets. Shaded areas indicate bootstrapped standard errors. Inset: corresponding action potentials (shaded area indicates standard deviation; vertical bar corresponds to 100 µV, duration: 2.5 ms).*

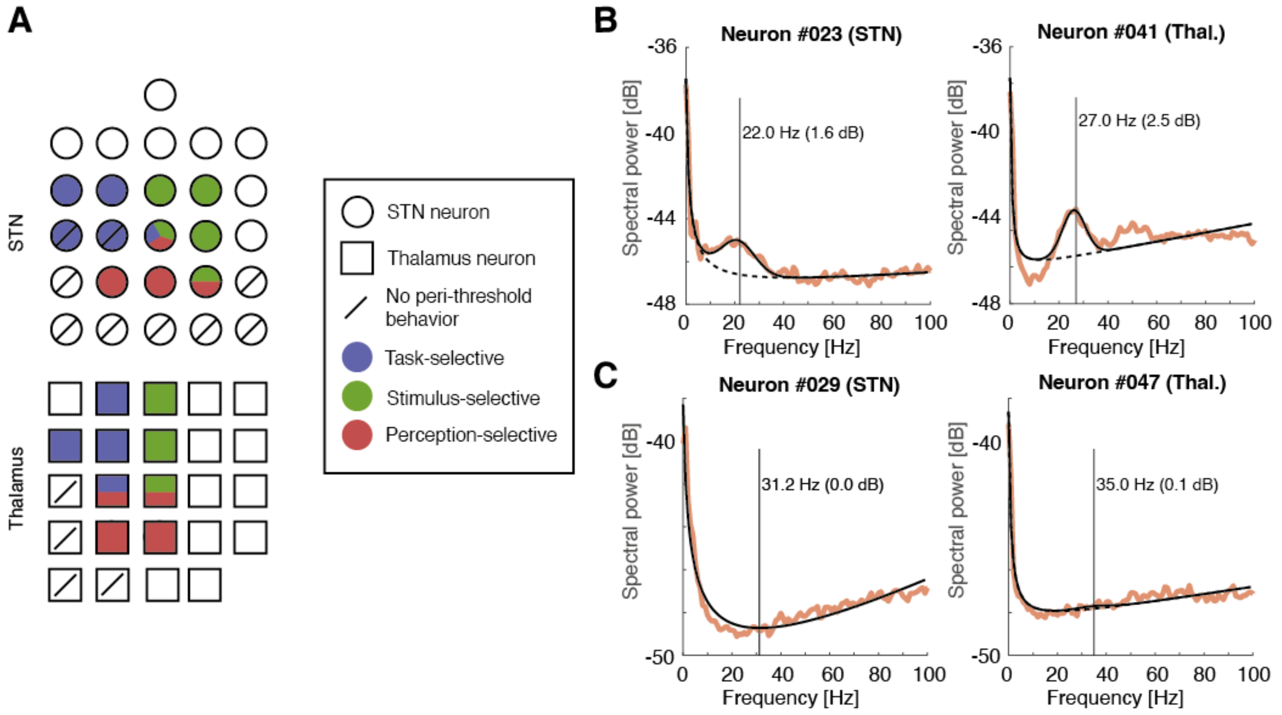


***Figure S7. A.*** *Distribution of STN (circles) and thalamus (square) neurons for task- (blue) stimulus- (green) and perception-selectivity (red). Neurons for which we did not obtain a sufficient number of trials for which behavior could be considered to be around the tactile perceptual threshold (see methods) are stricken by an oblique line.* ***B-C****. The resulting spectra (orange trace) and the resulting fit (black trace). To measure the beta component, we fitted the spectrum of the power spectral density of the spikes with 7 parameters: four to model the 1/f decay and 3 to model the bump in the spectrum: beta amplitude* (beta frequency + K*beta width), with K a Gaussian kernel of mean 0 and unit standard deviation [S1, S2]. We found no relationship between the selectivity of neurons and the frequency or amplitude of the beta component (p > 0.10). The spectrum without the beta component is plotted as a black dashed line.* ***B****. Neurons with the highest beta components in the STN (left) and thalamus (right).* ***C****. Neurons with the lowest beta components in the STN (left) and thalamus (right).*
